# Generation of cultured beef from bovine embryonic stem cells

**DOI:** 10.1101/2024.10.15.618593

**Authors:** Xue Rui, Zehan Li, Jiehuan Xu, Jianjun Dai, Xiaohui Zhang, Xiao Jin, Yang Liu

**Author notes:** These authors contributed equally: Xue Rui, Zehan Li. Corresponding authors: Yang Liu.

## Abstract

Bovine embryonic stem cells (bESCs) serve as an optimal source for generating muscle and fat tissues. However, systematic methodologies for creating cultured beef from bESCs have not yet been reported. Here, we successfully established a bESC line and formulated a specialized culture medium that supports cell pluripotency in both two-dimensional (2D) and three-dimensional (3D) cultures. Additionally, we have also developed serum-free and transgene-free protocols that efficiently differentiate bESCs into muscle and fat cells. These differentiated cells were validated at the transcriptional and protein levels. Furthermore, we devised a method using high moisture stretched plant-based protein scaffolds for incorporating differentiated muscle and fat cells derived from bESCs, producing an innovative plant-cell hybrid cultured beef. These advancements provide a solid foundation for future cultured beef production using pluripotent stem cells.

## Introduction

Cultured meat (also cultivated, cell-cultured, *in vitro* meat) uses tissue engineering to produce meat for food, offering a novel approach to meat production^1^. This innovation has attracted considerable interest for its capacity to mitigate challenges inherent in traditional livestock farming, such as environmental sustainability, animal welfare, public health, and food security^2^. Consequently, cultured meat development emerges as a viable solution to the problems plaguing conventional meat production, presenting an avenue for a more sustainable, ethical, and healthier food system. Nonetheless, substantial technological challenges must be addressed to fully harness its potential.

The advancement of cultured meat technology encounters numerous challenges, including the establishment of cell lines, development of cost-effective serum-free culture media, fabrication of textured scaffolds, replication of taste and nutritional profiles of commercial meats, and establishment of scalable, automated bioproduction processes^3^. The generation of high-quality cell lines is fundamental to cultured meat production. During cell line development, factors such as proliferation rate, differentiation efficiency into specific cell types like muscle and fat, and stability are critical. Current literature documents various cell lines for cultured meat, including primary adipose and muscle stem cells, alongside genetically modified organisms (GMO) and spontaneously immortalized cell lines. Primary stem cells offer promising differentiation capabilities but are limited by their proliferation capacity *in vitro*, necessitating regular animal sourcing^4–5^. Conversely, GMO^6^ and spontaneously immortalized cell lines^7^ facilitate continuous proliferation *in vitro* but encounter regulatory and consumer acceptance hurdles, impacting the perceived safety of cultured meat. Moreover, spontaneously immortalized cell lines have by far predominantly been explored in lower species such as chicken^7^ and fish^8^. Embryonic stem cells, not requiring genetic modification, exhibit indefinite and stable proliferation and versatile differentiation potential *in vitro*^9–10^. Establishing bESC lines has been a challenging task, with significant progress made only in recent years^11–13^. This advancement has paved the way for the application of ESC in cultured meat. However, the development of robust serum-free media for the expansion of ESC in feeder-free conditions, efficient protocols for inducing differentiation of bESCs into muscle and fat cells, as well as the development of cultured meat fabrication methods that are more aligned with commercial manufacturing scenarios, remain pressing issues that need to be addressed in the field^14–15^.

In this study, we established a bESC line and introduced a novel, customer-made culture medium, termed NBFR+ medium, designed to facilitate the stable passage of these cells under feeder-free conditions. Importantly, our findings indicate that this medium effectively maintains the pluripotency of bovine pluripotent stem cells in both 2D and 3D culture systems. Furthermore, we developed transgene-free and serum-free protocols that enable the differentiation of bESCs into muscle and fat cells, with the characteristics of these differentiated cells validated at the transcriptional and protein levels. In a final step, we combined muscle and fat cells derived from bESCs with textured plant-based protein chunks, thereby creating a novel plant-cell hybrid cultured beef.

## Materials

TrypLE Express (12604-013), DMEM/F-12 (11320033), Neurobasal (21103049), N-2 Supplement (17502001), B-27 Supplement (17504001), MEM NEAA (11140050), GlutaMAX (35050-061), Vitronectin (A14700), Pluronic™ F-68 (24040032), α-MEM (12571063), UltraPure Water (10977015), Activin A (120-14E), Sodium Pyruvate (11360070) and CD Lip concentration (12531018) were purchased from Thermo Fisher Scientific. Tyrode′s Solution (T1788-100ML), Triton™ X-100 (T8787), Bovine Serum Albumin (BSA, A1933), Mitomycin C (M5353), Dexamethasone (D4902), and Gelatin (G7041) were purchased from Sigma Aldrich. IWR-1-endo (S7086), Y-27632 (S6390), and Givinostat (GIVI, 732302-99-7) were purchased from Selleck. Penicillin-Streptomycin-Amphotericin B Solution (PSA, 4408584), Oil Red O Staining Kit (C0157S), Enhanced Cell Counting Kit-8 (C0042), Hoechst 33342 Staining Solution (C1029), Alkaline Phosphatase Assay Kit (P0321S), and HEPES (C0217) were purchased from Beyotime. HiPure Total RNA Mini Kit (R4111-02) and BODIPY (MX5403) were purchased from MGBio. 4% Paraformaldehyde Fix Solution (4% PFA, F8011) was purchased from Adamas Life. Fetal Bovine Serum (FBS, FSP500) was purchased from ExCell. Pronase (10165921001) was purchased from Roche. Low fatty acid BSA (4483930) was purchased from MP Biomedicals. Recombinant Human FGFb (C751) was purchased from Novoprotein. Dulbecco’s modified eagle medium (DMEM, SH30022.01) was purchased from Hyclone. Laminin (LN521-05) was purchased from BioLamina. TransScript® One-Step gDNA Removal and cDNA Synthesis SuperMix (AT311-03) was purchased from TransGen. ChamQ SYBR qPCR Master Mix (Q311-02) was purchased from Vazyme. SB-431542 (T1726) and 3-isobutyl-1-methylxanthine (IBMX, T1713) were purchased from TOPSCIENCE. Indomethacin (HY-14397) was purchased from MCE. Recombinant Human EGF (C029) was purchased from Novoprotein. TeSR™-E6 (05946) and FreSR-S™ (05859) were purchased from Stem Cell Technologies. Skeletal Muscle Cell Growth Medium-2 (SKGM, CC-3245) was purchased from LONZA. Defatty acid BSA (dBSA, S25762) was purchased from MedMol. MSC serum-free medium (S101-500/S102-025) was purchased from Baidi Biotechnology. Gluten from wheat, corn starch and soy protein isolate (SPI) were purchased from Feng Rui Biotechnology, China. Beet root powder was purchased from Tong Ren Tang, China. Sodium alginate was purchased from Feng Rui Biotechnology, China.

## Methods

### Isolation of MEF

Mouse embryos at E12.5 were carefully harvested from the uterus of pregnant mice. After the fetal head, limbs, tail, and internal organs were removed, the torsos were finely minced with sterilized surgical scissors until achieving uniform and small pieces (diameter<3 mm). These minced torsos were subsequently digested with 0.25% Trypsin-EDTA at 37°C for 10 minutes. The trypsinization process was terminated by the addition of MEF medium containing DMEM, 10% FBS and 1% PSA. The resulted cells were cultured in a 5% CO_2,_ 37°C incubator. MEF cells (P3) intended for use as a feeder layer were treated with 10 µg/ml mitomycin C for 3 hours. These treated MEFs could either be frozen and stored in liquid nitrogen or used as feeder cells by plating them in 0.2% gelatin-coated petri dishes or well plates at a density of 60,000 cells/cm^2^.

### bESCs establishment from embryos

Holstein embryos at E5-7 developmental stage were sourced from the Shanghai Academy of Agricultural Sciences. Thawed bovine blastocysts were carefully transferred to Tyrode’s acid solution using a mouth pipette. Subsequently, the blastocysts were transferred to Pronase drops and observed under a stereomicroscope (HSZ300), when most of the zona pellucida disappeared. The Pronase treatment (2-lasting 2-3 minutes, facilitated the removal of zona pellucida. Afterwards, the blastocysts were rinsed in NBFR medium to eliminate any residual Pronase and then plated onto mitotically inactivated MEF feeders. Blastocysts failed to adhere to the MEFs were gently pressed to the bottom of 12-well plate using a 22G needle for attachment. The growth of ESC colonies was monitored daily, with typically evident growth observed within 1-2 weeks.

### Culturing, passaging and freezing of bESCs

When reaching ∼70% confluency in the MEF feeder system, colonies were enzymatically detached using TrypLE. Afterwards, the cells were resuspended in NBFR medium containing 10 μM Y-27632, and passaged at a ratio of 1:16. The cells were then inoculated onto mitomycin C-treated MEF feeder cells and feed daily. The cell culture and passage protocol in E8+ (TeSR™-E6, 20 ng/ml FGFb, 2.5 uM IWR-1-endo, 1% PSA and 20 ng/ml Activin A) within the feeder system closely parallels that of NBFR medium. When reaching ∼80% confluency in the feeder-free system, the colonies were enzymatically detached using TrypLE, resuspended in feeder-free medium containing 10 μM Y-27632, and passaged at a ratio of 1:8 in 0.5 μg/cm^2^ Laminin-coated 6-well plates with daily feeding. Of note, the feeder-free medium comprised of 50% DMEM/F-12, 50% Neurobasal, 1% low fatty acid BSA, 20 ng/ml FGFb, 5 uM IWR-1-endo, 20 ng/ml Activin A, 1% N-2 Supplement, 2% B-27 Supplement, 1% MEM NEAA, 1% GlutaMAX, 50 ug/ml Vitamin C, 1 uM CHIR-99021, 0.25 mM Sodium Pyruvate, 0.3 uM WH-4-023, 1% PSA and 10 ng/ml LIF. For cryopreservation, cells were frozen using FreSR-S in liquid nitrogen, with each cryovial containing 1 million cells suspended in 1 ml freezing solution.

### Embryoid body (EB) differentiation

The bESCs were employed for EB formation. When reaching 70-80% confluency, the cells were detached using TryPLE, and seeded in ultralow-adhesion petri dishes (Thermo Fisher Scientific) containing feeder-free medium supplemented with 1 μM Rocki. Half of the medium was replaced with fresh DMEM medium containing 10% FBS every other day. After 7 days, the EBs were transferred to gelatin-coated plates and cultured in DMEM medium containing 10% FBS, with feeding every other day. On day 7, cells migrating from the EBs were harvested for RNA isolation and gene expression analysis.

### 3D suspension culture

The bESCs were detached with TryPLE, when reaching 70-80% confluency. Subsequently, the cells were seeded in a 50 ml Glass Spinner Flask (CytoNiche) in feeder-free medium containing 1% F68 at a concentration of 1×10^6^ cells/ml. Half of the medium was replaced daily. Samples were collected every two days for live/dead staining and cell counting to monitor cell viability and proliferation.

### Alkaline phosphatase staining

The bESCs were fixed with 4% PFA for 15 minutes at room temperature. The staining was conducted according to the Alkaline Phosphatase Assay Kit procedures. After the staining, the cells were immersed in PBS and imaged with a Nikon ECLIPSE Ts2 inverted microscope.

### Karyotype analysis

The bESCs were cultured in medium containing 75 ng/ml colcemid for 2 hours at 38.5°C. Afterwards, when reaching confluency, the cells were detached with TryPLE, and then treated with 0.075 M KCl at room temperature for 20 minutes. Following this, the cells were fixed with methanol/acetic acid (3:1, v/v), centrifuged at 1000 rpm for 5 minutes, transferred to slides and air-dried overnight. The chromosome number was determined from stained chromosome using 5% Giemsa for 10 minutes. A total of 50 cells in metaphase were used for counting. When a predominant number of cells exhibited an identical chromosome count, it was acknowledged as the modal chromosome number.

### Immunofluorescence staining

The cells were fixed with 4% PFA for 15 minutes, permeabilized with 0.1% Triton X-100 in PBS (0.1% PBST) for 15 minutes, and then blocked with 1% BSA in PBS for 1 hour at room temperature. Subsequently, cells were incubated overnight with primary antibodies listed in Table S1 in 0.1% PBST at 4 °C. The next day, cells were rinsed in DPBS and incubated with secondary antibodies (Table S1) at a dilution of 1:500 in 0.1% PBST for 1 hour at room temperature. Next, cells were rinsed in DPBS three times and stained with DAPI for 10 minutes. The stained cells were imaged using Nikon ECLIPSE Ts2 inverted microscope, Olympus CKX 53 inverted microscope, or Confocal microscope (Leica STELLARIS 5).

### Oil Red O Staining

The bESCs were fixed with 4% PFA for 15 minutes at room temperature, and then stained following the Oil Red O staining kit procedures. Finally, cells were rinsed in wash buffer twice, stored in PBS, and imaged using a Nikon Ts2 inverted microscope.

### Adipogenic differentiation of bESCs

Initially, bESCs were thawed and seeded in a matrix-coated 6-well plate at a density of 1×10^5^ cells/well in feeder-free medium supplemented with 10 μM Y-27632. When reaching 100% confluency, the cells were switched to adipogenic induction medium consisting of 50% Neurobasal, 50% DMEM/F12 medium, 1% N2, 2% B27, 3 μM CHIR-99021 and 10 μM SB-431542. During the first stage of differentiation, the cells were rinsed in DPBS prior to being cultured in neural ectoderm medium for up to 5 days, with feeding every other day. Subsequently, the cells were transitioned to commercial MSC serum-free medium for the second stage of differentiation, aiming for PAD formation after 4 days. When reaching ∼90% confluency, the PADs were switched to our self-developed medium for 5 days of lipid accumulation in a 5% CO_2,_ 38.5°C incubator, with feeding every other day. The cells were rinsed gently twice in DPBS to remove any residual MSC serum-free medium. During feeding, great care was taken to handle the cells gently to prevent lift-off of lipid-laden adipocytes. Tiny lipid droplets were observed within 2 days. Our self-developed differentiation medium consisting of 16 components, including DMEM/F12, dBSA, Hepes, and CD Lip concentration, while the conventional medium comprised of DMEM, 10% FBS, 1 μM dexamethasone, 0.5 mM IBMX, 0.2 mM indomethacin and 10 μM insulin.

### Myogenic differentiation of bESCs

Initially, bESCs were thawed and seeded in a matrix-coated 6-well plate at a density of 6×10^5^ cells/well in a 5% CO_2_, 38.5°C incubator. After 24 hours, the cells were switched to differentiation medium-1, consisting of E6 (as the basal medium supplemented with 10 μM CHIR99021) for 2 days. Following this, the cells were switched to differentiation medium-2, consisting of E6 supplemented with 100 nM GIVI for 5 days to complete the initial differentiation. Next, the cells were cultured in E6 for 7 days with daily feeding. The resulting MPCs exhibited sustained expansion capacity, as evidenced by their ability to undergo passage for up to 10 generations in SKGM medium. After reaching ∼90% confluency, the MPCs were switched to Neuro-2 medium, consisting of DMEM/F12 supplemented with1% ITS, 1% N2, and 1% GlutaMAX (differentiation medium) for 7 days with feeding every other day.

### qRT-PCR analysis

RNA was extracted and purified using the HiPure Total RNA Mini Kit according to the manufacturer’s instructions. The concentration and purity of RNA were assessed using Thermo Scientific NanoDrop Microvolume Spectrophotometers. Subsequently, cDNA was synthesized using the cDNA SuperMix and Thermal Cycler T100 following the manufacturer’s instructions. The primers (Table S2), SYBR FAST Universal 2× qPCR Master Mix, and LightCycler® 480 Instrument II were used for the analysis of gene expression. The transcription levels of genes were determined using the ΔΔCt method and normalized to GAPDH.

### RNA sequencing (RNA-seq) analysis

RNA was extracted from bESCs and bovine fibroblasts using HiPure Total RNA Mini Kit according to the manufacturer’s directions. Library preparation and RNA sequencing were performed by Shanghai Biochip Co., Ltd. Library construction was conducted using VAHTS Stranded mRNA-seq Library Prep Kit (NR602, Vazyme) and sequenced on Illumina HiSeqX with paired-end, 150 base pair (bp). Sequencing reads were mapped to the U.S. Department of Agriculture, Agricultural Research Service Bovine transcriptome (genome build ARS-UCD1.3) using hisat2 version 2.2.0, and expression levels of all genes were quantified using htseq-count version 0.13.5. htseq-count yield an expression matrix of inferred gene counts. Normalized reads count, PCA and differential expression analysis was performed using R package edgeR with default parameters.

### Graphic display of sequencing data

R studio (https://www.rstudio.com/) was used to perform principal component analysis (R package FactoMineR, factoextra, ggsci, ggpubr). Hierarchical clustering and heat maps were created by R package pheatmap.

### Manufacturing of cultured beef

The manufacturing process of cultured beef involved a two-step procedure. Initial, the edible plant-based protein scaffold was fabricated utilizing a modified mechanical elongation method^1^. The preparatory process for the scaffolds is depicted in Fig. S3, while the comprehensive information on the formulation of each component is provided in Tables S3 and S4. Adipocyte-like and myocyte-like cells underwent purification through washing in DPBS, followed by preservation at −20°C until their utilization. In the subsequent phase, cultured fat and muscle cells were introduced into the scaffold, constituting 20% of the total weight (with 10% attributed to the fat component and an additional 10% allocated to the muscle component), together with beet root powder, corn starch, and gluten. The resulting mixture was shaped as designed, such as a burger, and subsequently stored at 4°C for further characterization. Empty scaffold and commercial animal beef were employed as negative and positive controls, respectively.

### Texture profile analysis (TPA)

Texture Profile Analysis (TPA) tests were conducted employing a Universal TA Texture Analyzer (Shanghai Tengba Instrument Technology Co. Ltd., China) at room temperature. All samples were standardized to the same dimensions (25 mm in diameter and 5 mm in height) for consistency. A stainless plunger with a diameter of 36 mm was employed as the compressive probe. Consistent experimental conditions prevailed across all measurements. The pretest speed, test speed, and post-test speed were all calibrated to 1 mm/s based on the preliminary test results. The compression ratio was standardized at 30%, with a trigger force of 8 g and a 5-second interval between successive compressions. Upon completion of the second compression cycle, the probe returned to the trigger point and then to its initial position.

### Statistical analysis

Statistical analyses were performed using GraphPad Prism 9. The specific analyses utilized are detailed in the text and comprise Analysis of Variance (ANOVA) with Tukey’s post-hoc tests, t-tests with Welch’s correction, principal component analyses, and chi-square tests. Error bars and ± ranges denote standard deviations unless otherwise specified with SEM (standard error of the mean). The P value of 0.05 served as the threshold for statistical significance.

## Results

### Establishment of bESCs suitable for development of cultured meat

Embryonic stem cells, characterized by their infinite self-renewal capability and pluripotent differentiation potential, are considered ideal seed cells for the cultured meat. Our initial focus was on the feasibility of establishing bESC lines from the blastocysts of Chinese Holstein dairy cows.

Employing the NBFR culture system^16^, we dissolved the zona pellucida from intact embryos at embryonic day 6-7 and inoculated the isolated bovine blastocysts onto mouse embryonic fibroblast (MEF) feeder layers (Fig. 1a). Significant outgrowth was observed by the fourth day post inoculation. Subsequent passages involved removing trophoblast cells, resulting in densely cellular colonies with sharp edges. Totally, we successfully isolated 10 bESC lines from 18 Holstein cow embryos. These cell lines have been continuously passaged, now reaching over 50 generations. Further validation demonstrated that our bESCs exhibited the capability to be passaged as either individual cells or small clusters, with positive results in alkaline phosphatase assays (Fig. 1b). Throughout long-term passaging, bESCs maintained a stable doubling time of approximately 24 hours (Fig. 1c) and a normal karyotype (2n = 60) (Fig. 1d). Immunofluorescence staining confirmed the expression of pluripotency markers such as NANOG, SOX2, OCT4, and SSEA4 (Fig. 1e). Gene expression analysis via qPCR further verified the high expression and maintenance of pluripotency markers over time (Fig. 1f, g). To assess differentiation capabilities, we cultured embryonic bodies (EBs) in ultra-low attachment plates, which spontaneously differentiated into three germ layers, as evidenced by specific marker expressions (Fig. 1i, h). Principal component analysis (PCA) of RNA-seq data conclusively demonstrates the distinct differentiation between bESCs and bovine fibroblasts (bFBs) (Fig. 1j). Consistent with prior assumptions, RNA-seq confirms the expression of unique classical biomarkers in each cell type (Fig. 1k). We next evaluated the applicability of NBFR medium for feeder-free culture of the established Chinese Holstein dairy cow ESC. Culturing these cells without a feeder layer led to significant cell death, with surviving cells diverging from clonal growth and exhibiting signs of differentiation and reduced proliferation (Fig. 1l). Through iterative refinement, we developed a new culture medium, designated NBFR+, which notably supported dense clonal formation under feeder-free conditions (Fig. 1l).

**Fig. 1:**
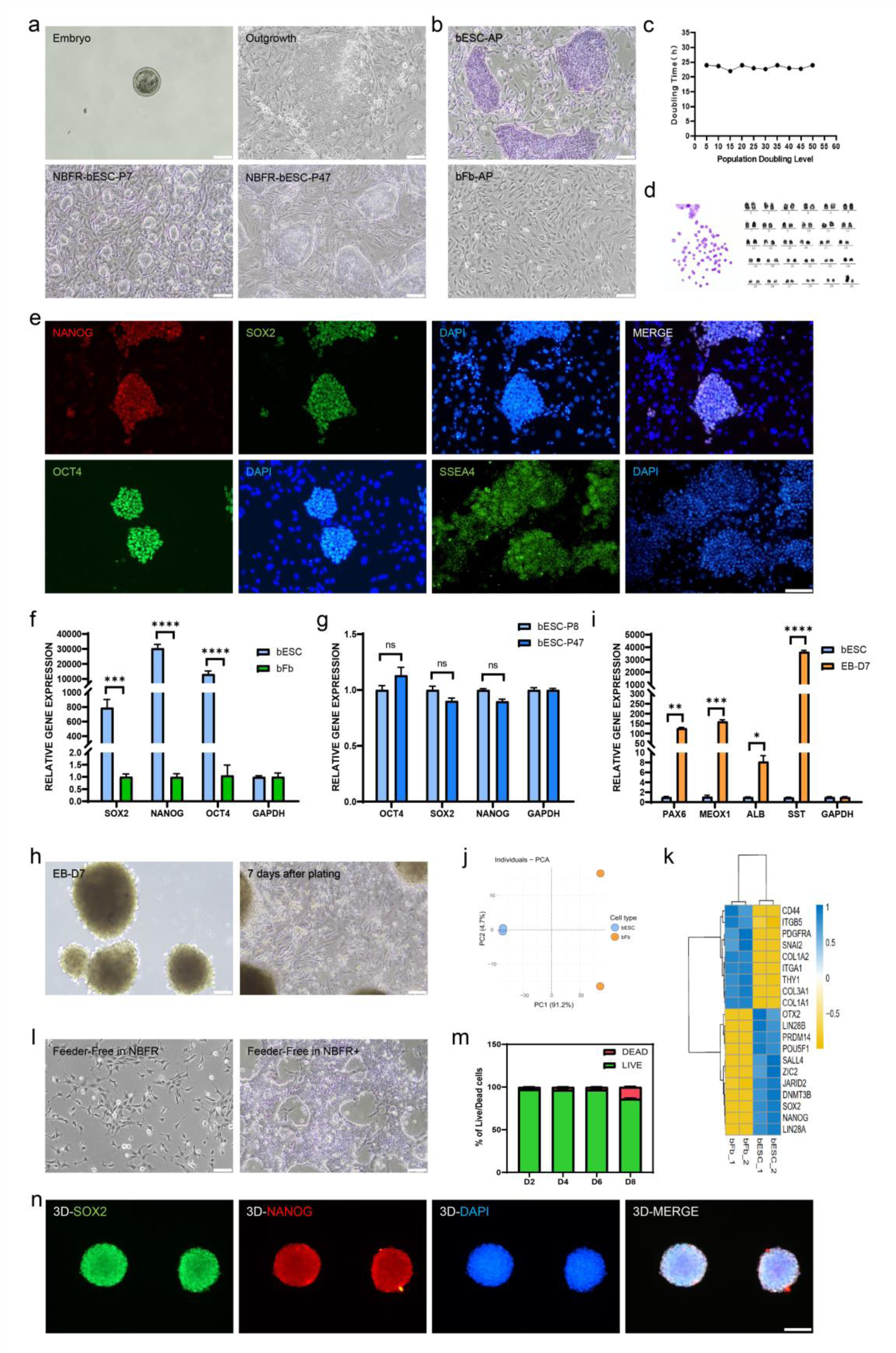
Establishment, characterization and culture medium optimization of bESCs. **a,** Representative bright field images of bESCs derived from the whole bovine blastocysts at various stages. These images represented a selected E5 embryo for bESC derivation, outgrowth in culture, typical colony morphology of bESCs at P7 and P47 when cultured in NBFR medium. **b,** Comparison of alkaline phosphatase (AP) activity of bESC and bFb. **c,** The doubling time of bESCs, stabilized at 23 ± 2 hours within 50 generations. **d,** Karyotype analysis of bESCs (P10), maintained normal chromosome numbers (2n = 60). **e,** Immunofluorescence staining of pluripotency markers (NANOG, SOX2, OCT4) and live staining of cell morphology marker (SSEA4) of bESCs. DAPI was used for nucleus staining. **f,** Comparison of gene expression related to pluripotency markers (SOX2, NANOG, OCT4) between bESC and bFb. **g,** Gene expression related to pluripotency of bESCs at P8 and P47. **h,** Gene expression of ectoderm markers (PAX6, SST), mesoderm maker (MEOX1) and endoderm marker (ALB) of spontaneous three germ layer differentiation for 7 days. **i,** EB formation after 3D suspension culture and cell morphology after adherent differentiation on day 7 day. **j,** Principal component analysis (PCA) plot of RNA-seq from bESC and bFb. **k,** Heatmap of differentially expressed gene of bESCs and bFb. **l,** Feeder-free bESCs cultured in NBFR medium. **m,** Percentage of live and dead bESCs on day 2, 4, 6 and 8 during 3D suspension culture, which were calculated from the of results of live/dead staining images. **n,** Immunofluorescence staining of pluripotency markers (SOX2, NANOG) of bESC spheroids after 3D suspension culture on day 8. DAPI was used for nucleus staining. P refers to passage. Scale bars equal 100 μm. Data are expressed as mean plus standard error of the mean (n=3). P values are labelled: ns (no significance) indicates p>0.05, *p < 0.05, **p < 0.01, ***p < 0.001, ****p < 0.0001.

Comprehensive analysis, including immunofluorescence staining and qPCR, verified the sustained expression of stemness genes in cells cultured with feeder-free NBFR+ system (Fig. S1). Encouraged by NBFR+’s efficacy in maintaining stem cells properties without feeder layers, we explored its utility in suspension cultures to facilitate large-scale 3D cell expansion, inoculating bESCs into a glass spinner flask. We continuously monitored the cell status over 8 days. We found suspension cultured bESCs formed spheroids, maintaining over 96% viability by day 6 (Fig. 1m, S2), while continuing to express pluripotent genes on day 8 (Fig. 1n). Overall, we successfully established the bESC line, which exhibited self-renewal capabilities and potential for multi-lineage differentiation. Moreover, we developed a new serum-free culture medium that support the maintenance of stemness under feeder-free culture conditions.

### A serum-free approach for efficient differentiation of bESCs into adipocytes

The aroma of meat originates from the oxidation of fats. Therefore, we explored whether bESCs could be efficiently differentiated into adipocytes. Previous research in humans has shown that the induction of the neuroectoderm can effectively differentiate pluripotent stem cells into mesenchymal stem cells^17^, which are adipogenic precursors with the capacity to generate fat. Thus, we hypothesized that the similar approach might guide the differentiation of bESCs into adipocytes.

To induce bESCs into adipogenic precursors, we initially added 10 μM SB431542 and 3 μM CHIR99021 to the culture medium to induce the formation of neuroectoderm. After 5 days, the cells exhibited a neural-like morphology and spread out from each other (Fig. 2a). qPCR analysis indicated an elevated expression of neuroectodermal markers such as P75 and ZIC1 post-induction (Fig. 2b). To further induce these neuroectodermal cells into adipogenic precursors, the cells were switched to a serum-free mesenchymal stem cell (MSC) medium. After 4 days, the cells adopted a spindle-shaped morphology (Fig. 2a). qPCR analysis revealed that cells at this stage expressed classic adipogenic precursor markers such as NEIF3K, CD90, and CD105 (Fig. 2c). Notably, these cells could proliferate in this serum-free medium, with markers such as ITGA5 exhibiting an increase from passage 0 to passage 2 (Fig. 2c). A classic adipocyte maturation medium was employed to further mature the adipocytes^18^. However, we found limited efficiency of this medium in differentiation of bovine cells to adipocytes (Fig. 2d). Consequently, we developed the adipogenic maturation serum-free medium (self-developed medium). We found adipogenic precursors derived from bESCs were efficiently differentiated into mature adipocytes within 5 days using our self-developed medium (Fig. 2d). qPCR further verified a notable augmentation in mature adipocyte markers in cells (Fig. 2e). Remarkably, adipocyte precursor cells cultured in our self-developed medium displayed visible lipid accumulation within 7 days (Fig. 2f). Moreover, combining the harvested fat cells with sodium alginate hygrogel resulted in a macroscale construct that resembled bovine adipose tissue (Fig. 2g). We next performed lipidomic analysis and obtained quantifiable comparison of cultured fat with native bovine subcutaneous fat tissues. Interestingly, cultured fat contained lower levels of palmitic acid (C16:0), and higher levels of oleic acid (C18:1) (Fig. 2h), compared to commercial beef fat, which has been reported to correlate positively with palatability^19^. Indeed, the proportion of monounsaturated fatty acids and polyunsaturated fatty acids increased, while saturated fatty acids reduced in cultured fat (Fig. 2i). Taken together, these results demonstrated that we were able to efficiently induce bESCs into mature fat cells with a serum-free and transgene-free approach.

**Fig. 2:**
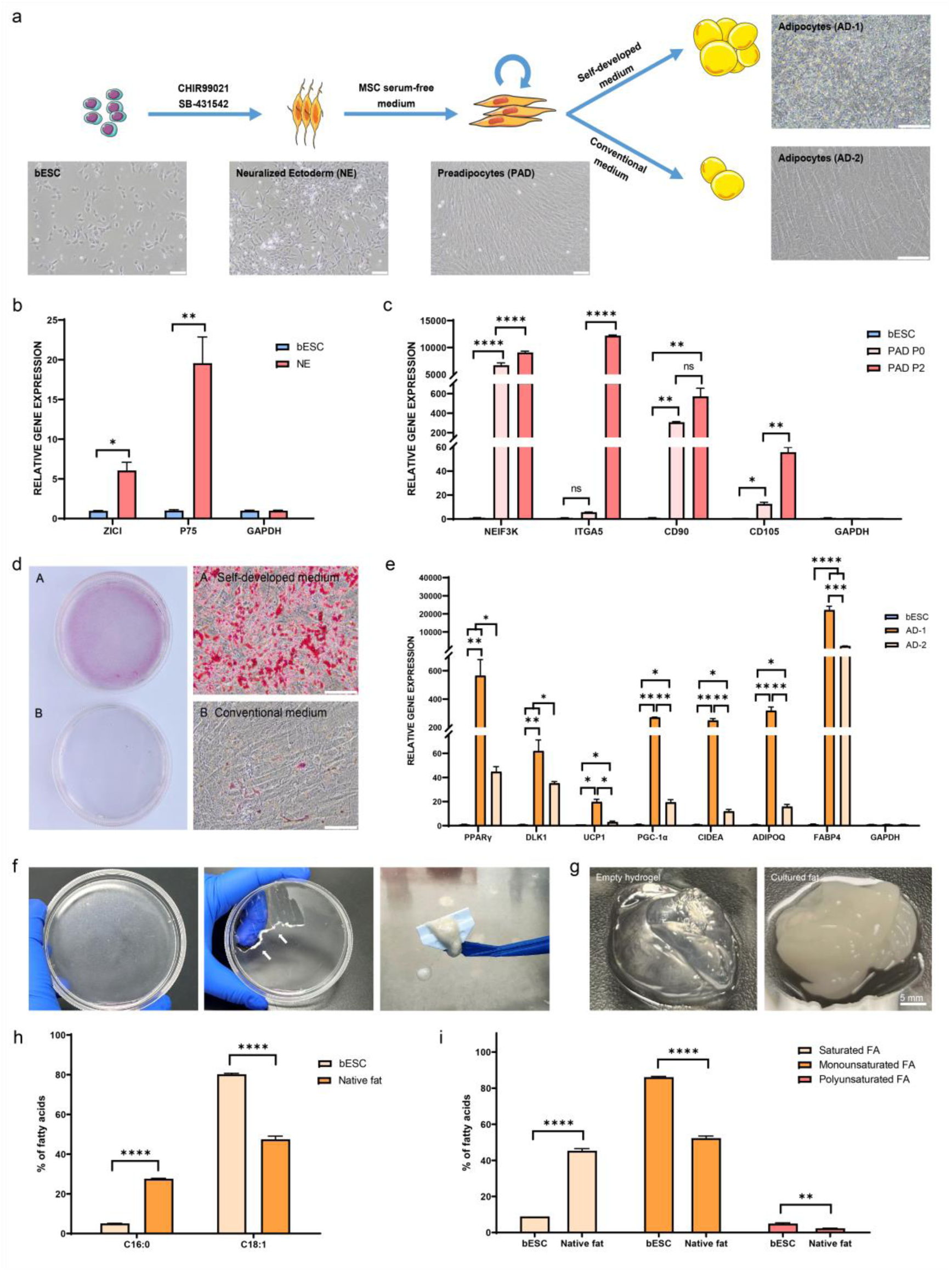
Evaluation and application of adipocyte differentiation in vitro. **a,** Schematic illustration of directional differentiation of bESCs to bovine adipocyte using our self-developed and conventional medium. **b,** Quantitative PCR (qPCR) analysis of neuroectoderm gene expression from bESCs and Neuralized Ectoderm (NE). **c,** Gene expression of preadipocytes (PAD, P0, P2) and bESCs after post-adipogenic induction. **d,** Oil red O staining of adipocytes cultured with self-developed (A) and conventional (B) medium for 5 days. **e,** Gene expression of adipocyte 1 (AD1) cultured with self-developed medium compared to adipocyte 2 (AD2) cultured with conventional medium. **f,** Representative bright filed images of accumulated adipocytes, rendered opaque on the bottom of the petri dish (left), with white arrows indicating the scraped adipocytes (middle), and mechanically collected adipocytes using a cell scraper (right) after 7 days of differentiation. **g,** Formation of cultured fat by combining adipocytes with empty alginate hydrogel. 1.5 (w/v) % alginate solution was prepared by adding alginate powder to DI water, fully hydrated at 50°C with continuously stirring for 1-2 hours, and centrifuged to remove any impurities. The scraped adipocytes were mixed with 1.5 (w/v) % alginate at a 1:1 (volumetric) ratio. For cultured bovine fat, 50 mM calcium chloride was mixed with alginate/adipocytes at a 1:1 (volumetric) ratio to induce the gelation. **h,** Comparison of percentages of fatty acid species between bESCs after 7 days of differentiation and native fat tissue. **i,** Comparison of percentages of saturated, monounsaturated and polyunsaturated fatty acids between bESCs after 7 days of differentiation and commercial beef fat tissue. P refers to passage. Scale bars equal 100 μm. Data are expressed as mean plus standard error of the mean (n=3). P values are labelled: ns (no significance) indicates p>0.05, *p < 0.05, **p < 0.01, ***p < 0.001, ****p < 0.0001.

### Direct differentiation of bESCs into skeletal muscle cells

Muscle cells, alongside fat cells, constitute another critically important cell type in meat. Leveraging the pluripotent differentiation potential of bESC, we investigated whether bESCs could be efficiently differentiated into muscle cells. We initially used a serum-free medium containing 10 μM CHIR99021 to direct bESC towards mesodermal differentiation (Fig. 3a). After 2 days, the cells underwent a significant epithelial-to-mesenchymal transition (Fig. 3a). The mesodermal marker T and early muscle cell marker PAX3 were highly expressed in the cells at this differentiation stage, while pluripotency markers such as OCT4 and SOX2 were significantly downregulated (Fig. 3b). To further induce muscle progenitor cell (MPC) formation, we subsequently treated the cells with 100 nM Givinostat (a histone deacetylase inhibitor) in E6 medium for 5 days, followed by a week in basal E6 medium, leading to elongated, spindle-shaped cells (Fig. 3a). The MPCs expressed significantly higher levels of myogenic transcription factors, including MYOG and MYOD, compared to bESCs (Fig. 3c). Culturing MPCs in SKGM muscle stem cell expansion medium (LONZA) allowed for further propagation, partially maintaining muscle stem cell marker expression (Fig. 3d). The MPCs were incubated in N2 medium, led to more elongated and aligned cells after one week. We found increased expression of mature myogenic markers such as CAV3, MYHC, and MYOSIN (Fig3, e, f). Remarkably, Myoglobin (MB) gene expression significantly increased in myotubes (Fig. 3e), which is in contrast with prior studies that suggested limited MB expression in myotubes derived from adult muscle stem cells^20^. Myoglobin is significant in the color and flavor of meat underscores the potential advantages of using muscle cells differentiated from bESCs. These results present a novel method for the efficient differentiation of bESCs into muscle cells.

**Fig. 3:**
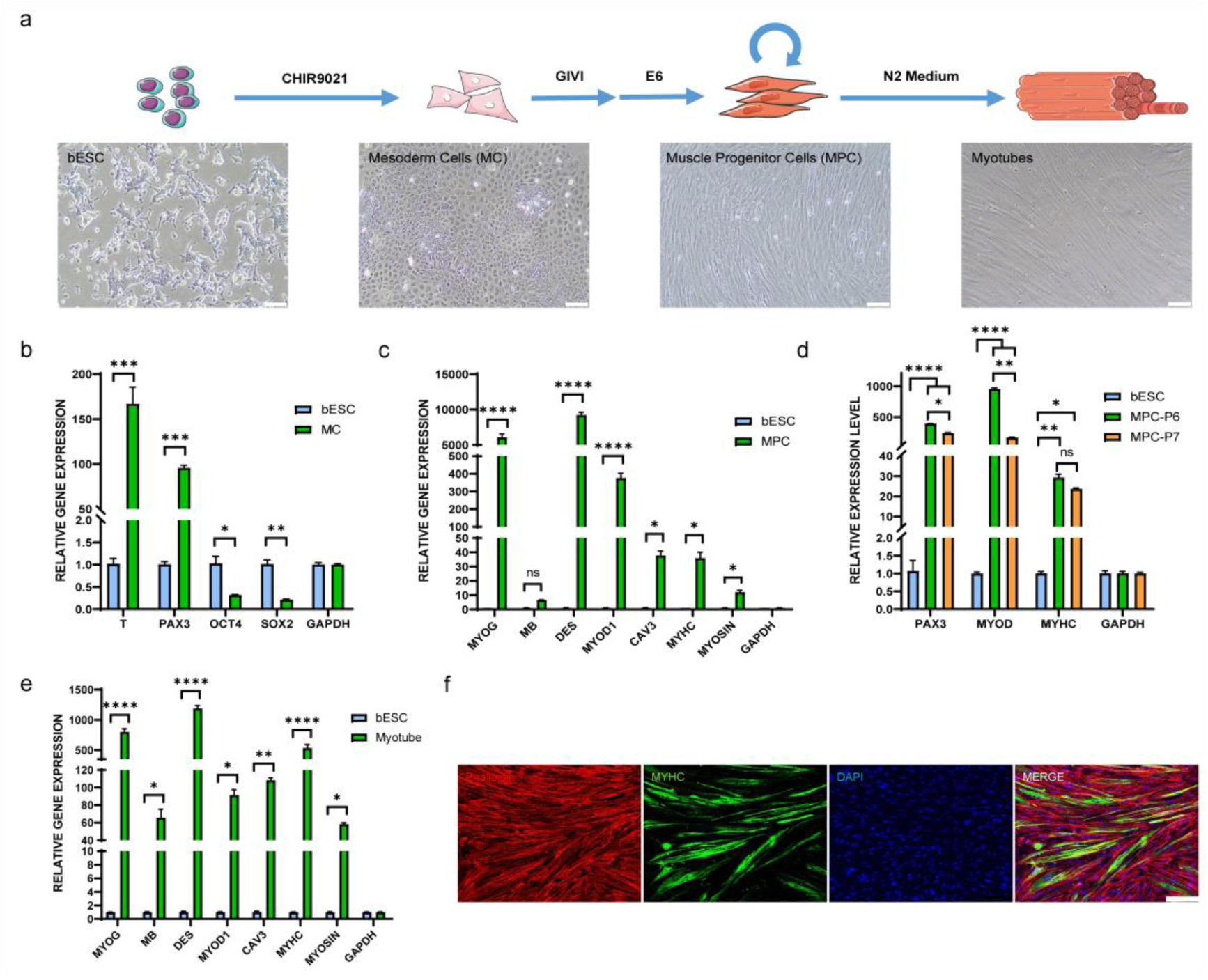
Evaluation of myocyte differentiation in vitro. **a,** Schematic illustration of directional differentiation of bESCs to bovine myocytes and myotubes, along with cell morphology at various stages. **b,** Gene expression of mesoderm markers (T, PAX3) and pluripotency markers (OCT4, SOX2) in myocytes compared to bESCs. **c,** Analysis of myogenesis markers (MYOG, MB, MYOD1, CAV3, MYHC, MYOSIN) expression in MPC (P6) compared to bESCs. **d,** Analysis of myogenesis markers (PAX3, MYOD, MYHC) expression in MPC (P6, P7) compared to bESCs. **e,** Analysis of myogenesis markers (MYOG, MB, DES, MYOD1, CAV3, MYHC, MYOSIN) expression in myotubes compared to bESCs. **f,** Immunofluorescence staining of myotubes (MF20, green), cellular actin (Phalloidin, red) and nuclei (DAPI, blue) from MPC after differentiation for 7 days. P refers to passage. Scale bars equal 100 μm. Data are expressed as mean plus standard error of the mean (n=3). P values are labelled: ns (no significance) indicates p >0.05, *p < 0.05, **p < 0.01, ***p < 0.001, ****p < 0.0001.

### Engineered cultured meat by the assembly of bESC derived fat and muscle cells with plant-based edible scaffolds

Another challenge in cultured meat is how to integrate scaffolding technology with cultured cells and closely resembles conventional meat in appearance, nutrition, and taste. Therefore, we developed a method for cultured beef in hamburger patties, integrating muscle and fat cells differentiated from bESCs with edible scaffolds containing gluten and soy protein isolate.

We first directed the differentiation of bESCs into muscle and fat cells (Fig. 4a). In scaffold preparation, soy protein isolate and gluten were processed into scaffolds with specific textural characteristics through mechanical stretching methods to better simulate the texture of meat (Fig. S3). Cultured muscle and fat cells were then thoroughly mixed with the structural protein scaffolds. Optical images of both raw and fried cultured beef demonstrate that the meticulous blending and molding of these components yield a composition closely resembling the sensory attributes and appearances of commercial beef (Fig. 4b). To further investigate the nutritional value of cultured beef, we conducted of its amino acid composition and content. All samples manifested the presence of nine essential amino acids, as anticipated with the inclusion of cultured cells and soy protein (Fig. 4c & S4). Significantly, the cultured beef exhibited higher contents of Valine, Leucine, Isoleucine, and Phenylalanine compared to animal beef in these amino acids (Fig. 4c). Subsequently, the texture profile analysis was conducted to investigate the textural properties of cultured beef, empty scaffold, and commercial beef (Fig. 4d). The values for chewiness and hardness of the empty scaffold significantly surpassed both cultured and commercial beef, possibly attributed to its textured structure and preserved intrinsic properties. In contrast to the empty scaffold, cultured beef demonstrated mechanical properties more closely resembling those of commercial beef, signifying a heighted similarity in texture between cultured and commercial beef. Overall, we successfully created a novel plant-cell hybrid cultured beef through combining muscle and fat cells derived from bESCs with textured scaffolds.

**Fig 4:**
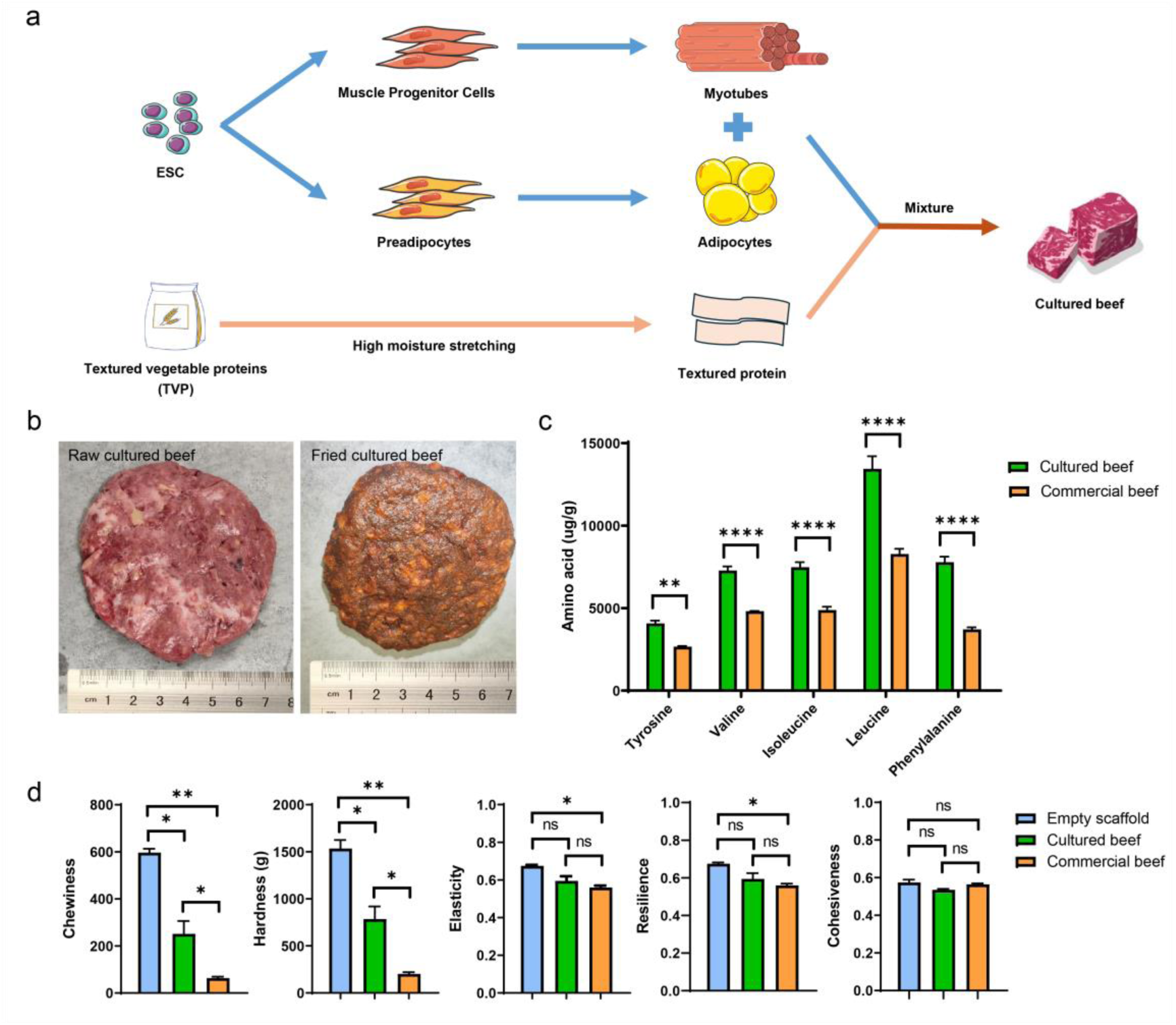
Manufacturing of cultured beef. **a,** A schematic illustration of the directional differentiation of bESCs to bovine adipocytes and myocytes, and subsequently combined with plant-based edible scaffolds. The edible scaffolds were sourced from soy protein isolate and gluten from wheat, underwent a systematic blending and hydration procedure, followed by meticulous mechanical elongation. Simultaneously, cells underwent cultivation and expansion processes to augment the overall biomass. The resultant cultured cells were seamlessly incorporated into the scaffold, resulted in cultured beef. **b,** Comparison of raw (left) and fried (right) cultured beef. **c,** Comparison of Tyrosine, Valine, Isoleucine, Leucine and Phenylalanine contents of cultured and commercial beef. **d,** Texture profile analysis of empty scaffold, cultured beef, and commercial beef. P values are labelled: ns (no significance) indicates p>0.05, *p < 0.05, **p < 0.01, ***p < 0.001, ****p < 0.0001.

## Discussion

The production of cultured meat necessitates a reliable supply of muscle and fat cells. Currently employed cell lines include primary, genetically modified, and spontaneously immortalized cell lines, each facing significant challenges. Primary cells require frequent animal slaughter for tissue acquisition due to limited proliferative ability, contradicting cultured meat’s ethical premise. Both genetically modified and spontaneously immortalized cell lines present food safety and regulatory hurdles and inadequate differentiation into muscle and fat cells. Spontaneously immortalized cell lines depend on selecting spontaneously mutated cells, a process feasible in lower species like fish and chicken but improbable in large mammals such as cattle^21^. Recently, induced pluripotent stem cells (iPSCs) have emerged as a promising technology in regenerative medicine, although their reprogramming involves exogenous gene transfection, prompting safety concerns in food production. Advances in non-integrating reprogramming technologies such as mRNA show potential in positioning iPSCs as a vital source for cultured meat from livestock cells. Our work utilized non-genetically modified bESCs derived from fertilized and *in vivo*-developed blastocysts. We found these cells can be cultured in a serum-free and feeder-free culture system, demonstrating rapid and stable proliferation, as well as maintaining self-renewal and pluripotency across continuous passages. Cultured meat technology aims to leverage suspension culture methods for enhanced cell proliferation and differentiation. Our bESC lines sustain pluripotency in a microcarrier-free 3D suspension culture, highlighting the potential of augmenting suspension culture scale and density through biotechnological advancements. Research on human pluripotent stem cells demonstrate that modifications in culture apparatus^22^, continuous perfusion techniques^23^, or thermoreversible hydrogels^24^ significantly increase the cell density.

Fat and muscle cells are pivotal to the cultured meat, necessitating efficient differentiation from seed cells. Challenges in differentiating primary adult stem cells include prolonged differentiation cycles^25^ and diminished capacity with increased passage numbers^4^. Previous studies on differentiation of pluripotent stem cells focus on human and mouse models, adopting two primary strategies: one is a step-by-step induction of pluripotent stem cells into specific cell types via defined signaling pathways^26–27^, plagued by lengthy cycles, complex steps, and partial dependence on animal serum; the other one employs genetic modification to hasten differentiation^28–29^, raising food safety and regulatory concerns. Our work developed a rapid, serum-free, and transgene-free protocol for differentiating bESCs into muscle and fat cells. We utilized a novel pathway inducing preadipocytes from the neuroectoderm, capable of *in vitro* proliferation and further rapid maturation using our self-developed adipogenesis medium. This method yielded mature adipocytes with a higher unsaturated fatty acid content compared to subcutaneous fat in cattle. Thus, our findings highlight the health advantages of unsaturated fats, particularly monounsaturated (MUFAs) and polyunsaturated fatty acids (PUFAs), known for their antioxidant and anticarcinogenic properties, while saturated fatty acids (SFAs) are associated with cardiovascular risks^30^. We established a more straightforward and efficient myotube differentiation protocol. qPCR verified the myoglobin expression in myotubes, distinguishing them from those derived from adult muscle stem cells^20^. Myoglobin is significant for beef’s color, taste, and nutritional value^31^. Further work will focus on transitioning these differentiation techniques from 2D cultures to 3D suspension systems, a method already explored in human pluripotent stem cell research for differentiation into various cell types, including cardiomyocytes^32^, hepatocytes^33^, endothelial cells^34^ and MSC^35^.

Plant-based proteins have received considerable attention in the food industry in respond to the increasing consumer preferences for ethical and eco-friendly alternatives to animal proteins^36^. Ideally, scaffolds derived from plant-based proteins should be cost-effective, thermally stable during cooking, non-toxic, and non-allergenic, exemplified by soy, pea, chickpea, zein, and wheat gluten. However, these individual plant-based proteins are often considered nutritionally incomplete, lacking some essential amino acids necessary for human consumption^37^. Consequently, a combination of proteins is necessary to achieve a balanced amino acid profile. To address the nutritional deficiencies, a blend of diverse plant-based proteins and incorporation of alternative proteins, such as cultured cells, in hybrid models, are being considered. Furthermore, hybrid meats, incorporating cultured cells, exhibit lower content of saturated fat and cholesterol along with fewer calories while maintaining protein richness compared to commercial meats. Various cell/scaffolding systems have been explored for hybrid meat production^38–40^. For example, the utilization of textured soy protein, an edible plant-based protein, in the cultivation of bovine satellite cells has demonstrated potential for high-quality cultured meat production^39^. However, these methods present challenges due to their complexity, cost, time consumption, and stringent scaffold requirements, restricting the scalability of cultured meat. In contrast, our approach involving the separate culturing of fat and muscle cells without scaffold involvement in the culture media, offering advantages in taste, sensory experience, texture, health, convenience, and scalability compared to methods involving scaffolds during culturing.

Cultured beef production requires a consistent cell line supply, with bESCs emerging as a promising source due to their limitless replicative capacity. The persistent pluripotency and genetic stability of our bESC lines form a crucial foundation for the consistent replication of cultured meat production. This not only ensures reproducibility but also eliminates the need for repeated animal biopsies, reducing costs and enhancing food safety. Additionally, the easy transportability of bESC vials minimizes logistical challenges and lowers the carbon footprint, marking a substantial step toward bolstering the sustainability of our cultured meat production process. Notably, cultured beef demonstrates cost competitiveness against commercial beef by addressing primary cost drivers, such as growth factors^41^, basal media^42^ and recombinant proteins^43^. For instance, Specht suggested aligning growth factor production with established enzyme production practices for optimization^44^. Integrating hybrid meat, combining plant-based ingredients with cultured cells, offers a near-term opportunity for cost reduction and a more authentic meat-eating experience. Moreover, technological advancements, including specialized cultivators, automated meat production, and improved growth factor production methods, show promise in substantially decreasing the cost of cultured meat^45^. The utilization of perfusion bioreactors is notable for generating high-density cultures, facilitating continuous cell growth. These insights underscore the economic significance of addressing cost drivers and optimizing production processes in cultured beef.

In this study, we elucidate the strategic methodologies to leverage the indefinite propagation and pluripotent differentiation attributes inherent in bESCs. Our objective is to establish an efficient and resilient production process for the generation of high-quality cultured meat products. Through comprehensive visual and sensory analyses, our findings reveal striking parallels between cultured and commercial beef (Fig. 4c). We have demonstrated the potential of cultured fat and muscle cells in serving as viable substitutes for commercial meat sources. This work represents a pioneering initiative poised to elevate the role of bESCs in the cultured meat production.

## Acknowledgements

We thank C Future for funding this work. Further, we thank Jiapei Wang, Xiaohong Chen and Mingqing Xu for technical support; Zhe Chen for concept discussion.

## Abbreviations

bESCs: bovine embryonic stem cells
2D: two-dimensional
3D: three-dimensional
GMO: genetically modified organisms
EB: embryoid body
SPI: soy protein isolate
TPA: texture profile analysis
ANOVA: analysis of variance
SEM: standard error of the mean
MEF: mouse embryonic fibroblast
PCA: principal component analysis
bFBs: bovine fibroblasts
MSC: mesenchymal stem cell
PAD: preadipocytes
AD1: adipocytes cultured with self-developed medium
AD2: adipocytes cultured with conventional medium
NE: neutralized ectoderm
MPC: muscle progenitor cell
MB: myoglobin
iPSCs: induced pluripotent stem cells
MUFAs: monounsaturated fatty acids
PUFAs: polyunsaturated fatty acids
SFAs: saturated fatty acids

## Author contributions

Xue Rui, Zehan Li, Jiehuan Xu, Xiaohui Zhang, and Xiao Jin performed experiments. Jianjun Dai donated the blastocysts of cattle. Xue Rui, Zehan Li, and Yang Liu drafted the manuscript. Yang Liu funded and supervised the entire work.

## Data Availability Statement

The datasets generated during and/or analyzed during the current study are available from the corresponding author on reasonable request.

## Ethics and guidelines statement

All procedures in the manuscript were reviewed in advance by the Laboratory Animal Ethics Committee of Shanghai Academy of Agricultural Sciences and also met the guidelines of the National Institutes of Health Guide for the Care and Use of Laboratory Animals.

---

**Figure S1.**
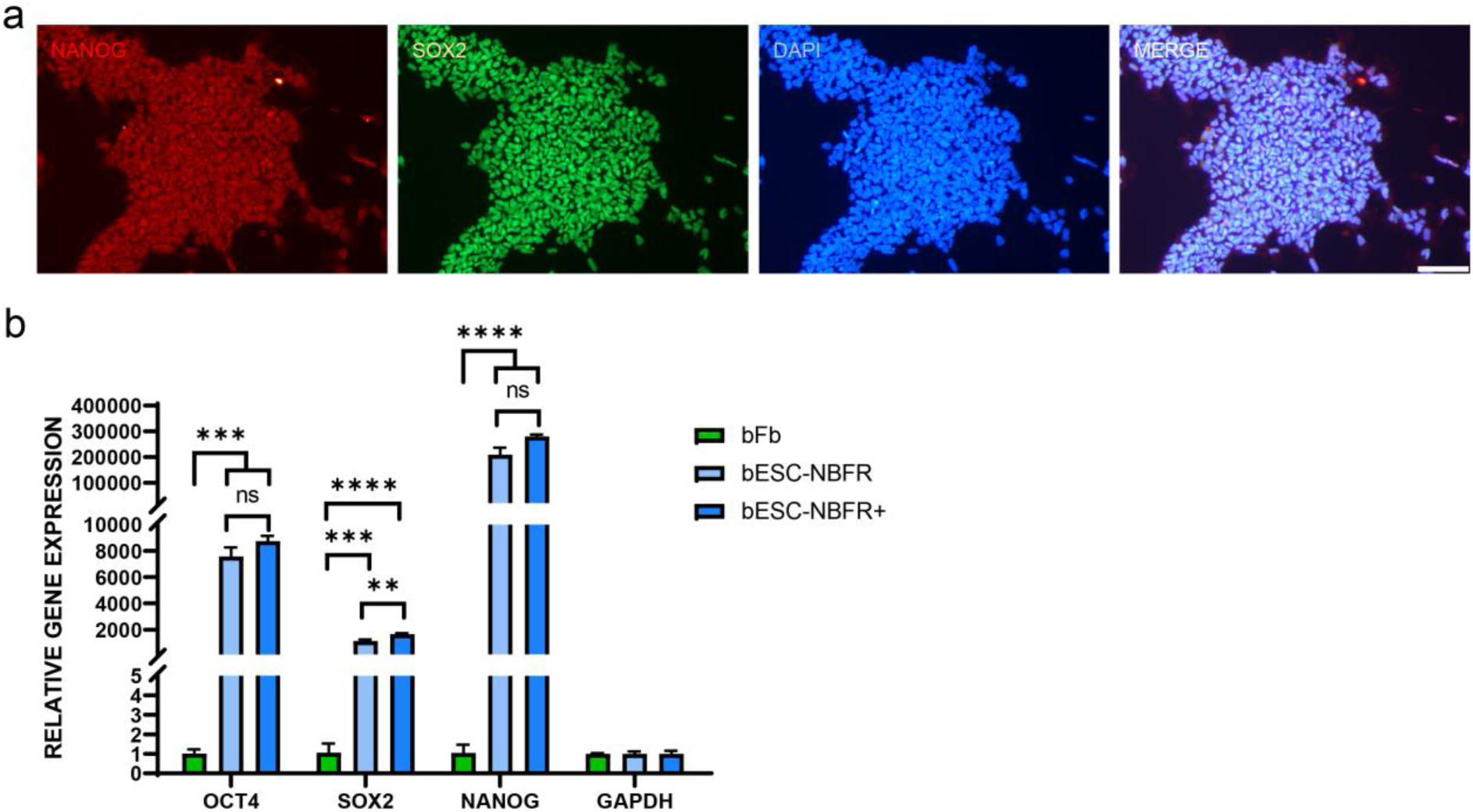
**a,** Immunofluorescence staining of pluripotency markers (NANOG, SOX2) on feeder-free bESCs, cultured in NBFR+ medium. Scale bars equal 100 μm. **b,** Gene expression analysis of pluripotency markers (OCT4, SOX2, NANOG) in bESC on feeder in NBFR, and feeder-free bESCs in NBFR+, compared to negative control bFb. (n=3) Data are expressed as mean plus standard error of the mean. P values are labelled: ns (no significance) indicates p>0.05, *p < 0.05, **p < 0.01, ***p < 0.001, ****p < 0.0001.

**Figure S2.**
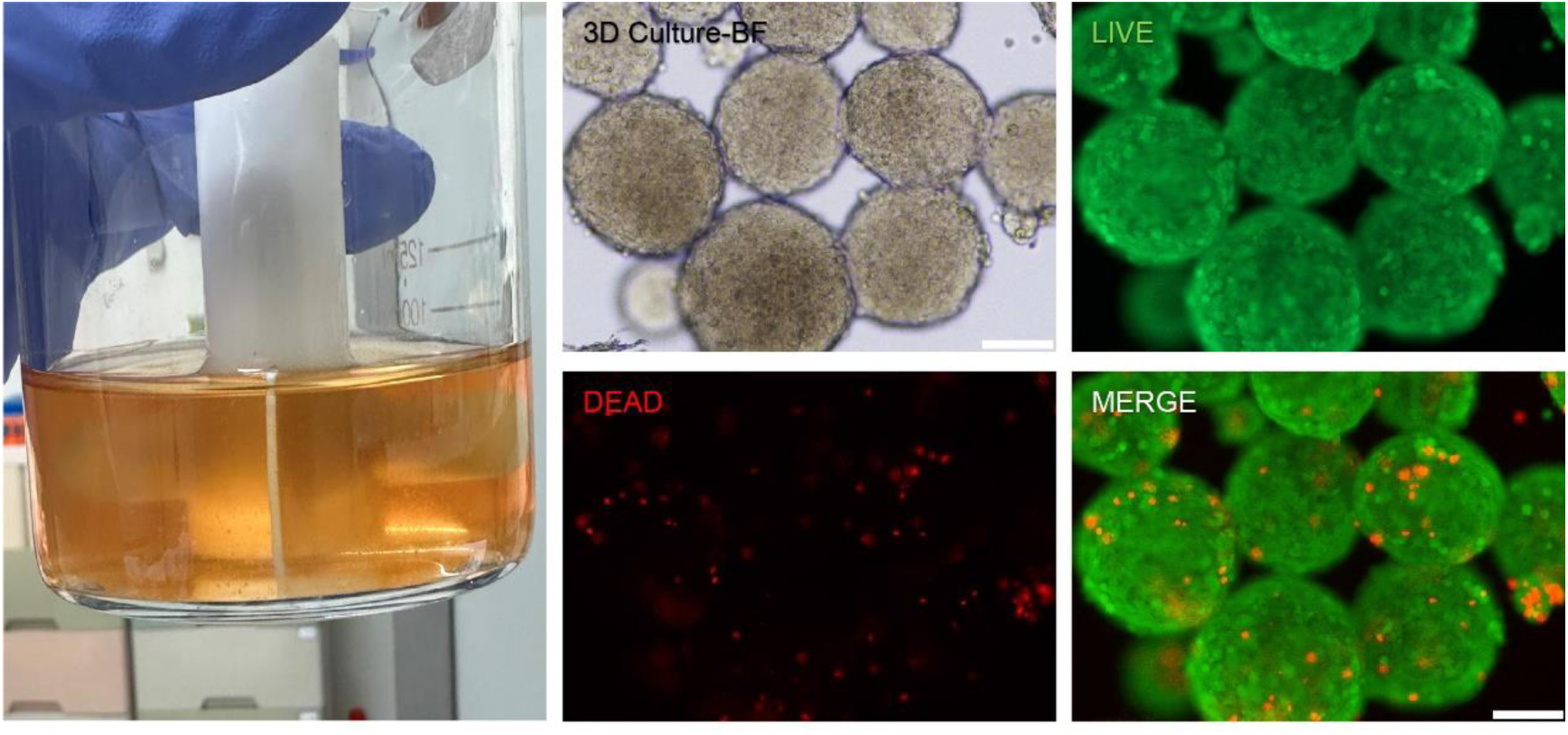
3D suspension culture of bESC (left). bESC-spheroids cultured in a glass spinner flask with NBFR+ medium and 1% F68. Bright field images of spheroids after 8 days of suspension culture and their live/dead rate (right). Scale bars 100 μm.

**Figure S3.**
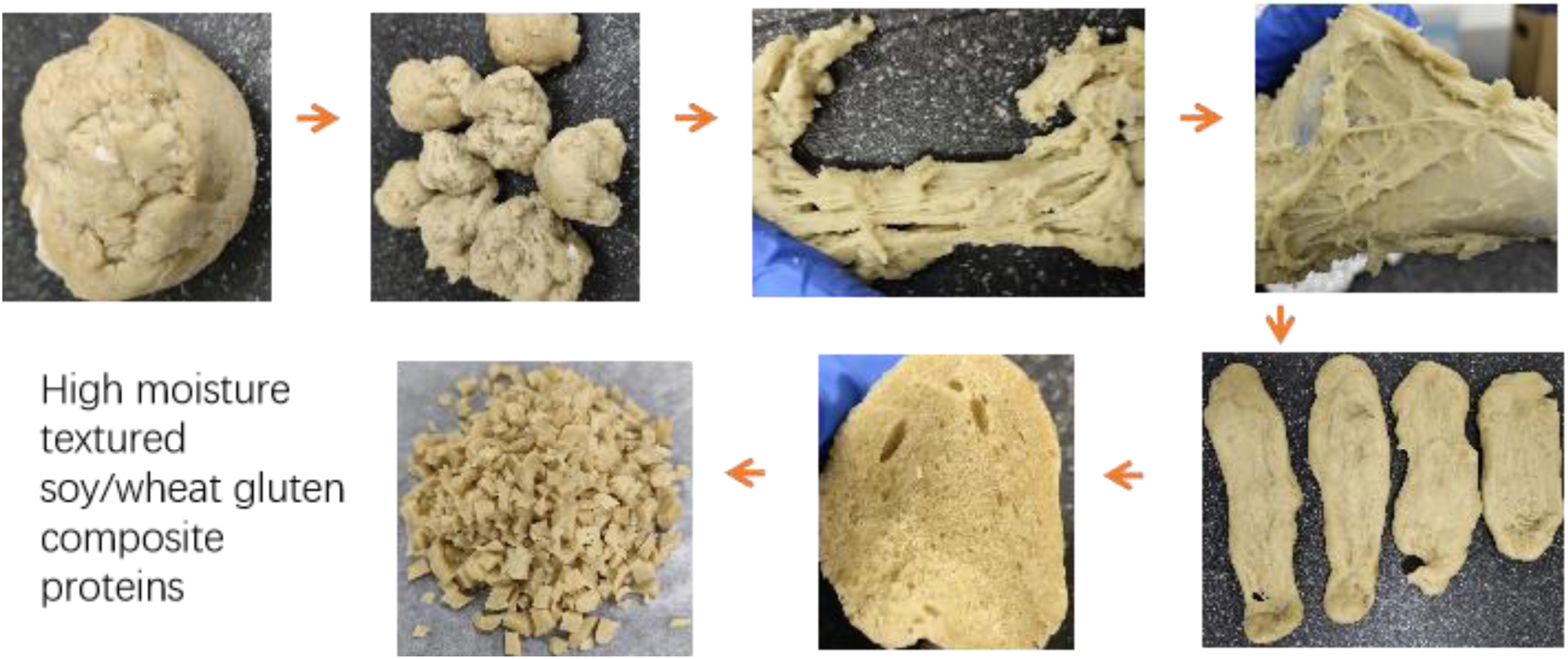
A schematic illustration elucidating the mechanical elongation method employed in the fabrication of textured protein scaffolds as described by Chiang et al.^1^ with modification. Briefly, the formulation process involved the meticulous hand-mixing of soy, gluten, and deionized water at room temperature to achieve a homogeneous dough. Subsequently, the dough underwent a detailed incubation process in a water bath at 60°C for one hour, ensuring comprehensive hydration of the gluten component. Following this hydration phase, the dough was methodically torn into pieces. It was postulated that an additional sequence of stretching and pulling would align the protein network, thereby facilitating protein agglomeration and engendering the formation of robust networks. Conforming to this hypothesis, the doughs were subjected to elongation, reaching twice their initial length, and were then meticulously folded to attain the ultimate dough structure. Each resulting dough was carefully encased in baking paper and subsequently covered with a layer of aluminum foil before undergoing a 30-minute steaming procedure. The dough was then methodically sectioned into diminutive pieces and judiciously stored at 4°C for prospective utilization.

**Figure S4.**
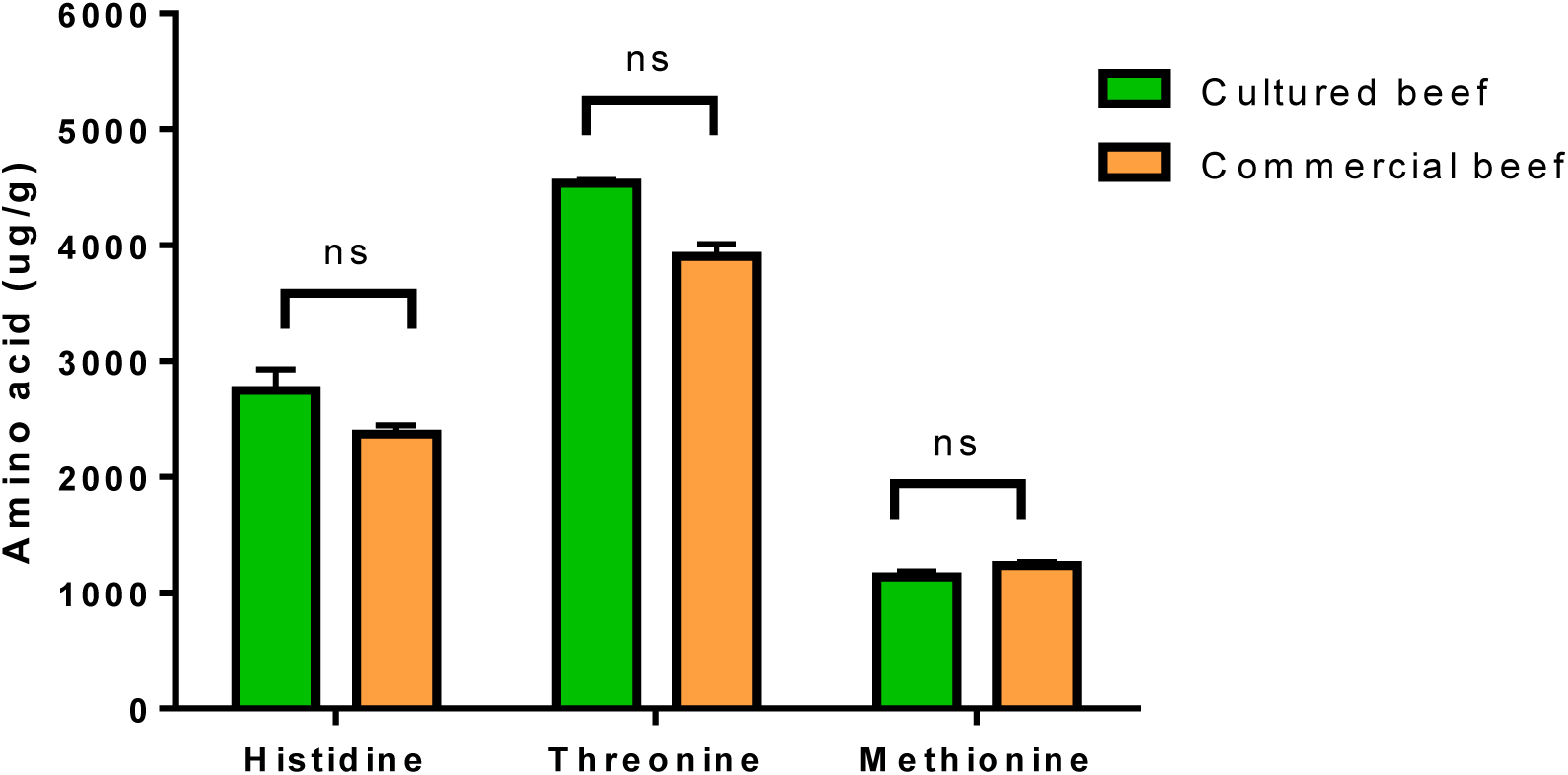
The amino acid analysis was conducted using the ultra-high-performance liquid chromatography (UHPLC, QE) Thermo, USA) coupled with a high-resolution orbitrap mass spectrometer. The samples underwent extraction at room temperature for 1 hour in the presence of 0.5 ml of 0.1 M hydrochloric acid (HCl), followed by centrifugation at 12,000 rpm for 10 minutes, with subsequent collection of the supernatant. Next, 10 μl of the diluted supernatant was subjected to ultra-high-performance liquid chromatography (UPLC) analysis. The chromatographic column was a Waters BEH C18 (50 x 2.1 mm, 1.7 μm), maintained at 55°C. The injection volume was set at 1 μl, and the flow rate was maintained at 0.5 ml/min. Mobile phases comprised ultrapure water (phase A) and acetonitrile (phase B) containing 0.1% formic acid. The elution gradient proceeded as follows: 0 min 95% A, 5.5 min 90% A, 7.5 min 75% A, 8 min 40% A, 8.5 min 95% A, and 13 min 95% A. P values are labelled: ns (no significance).

**Table S1.**
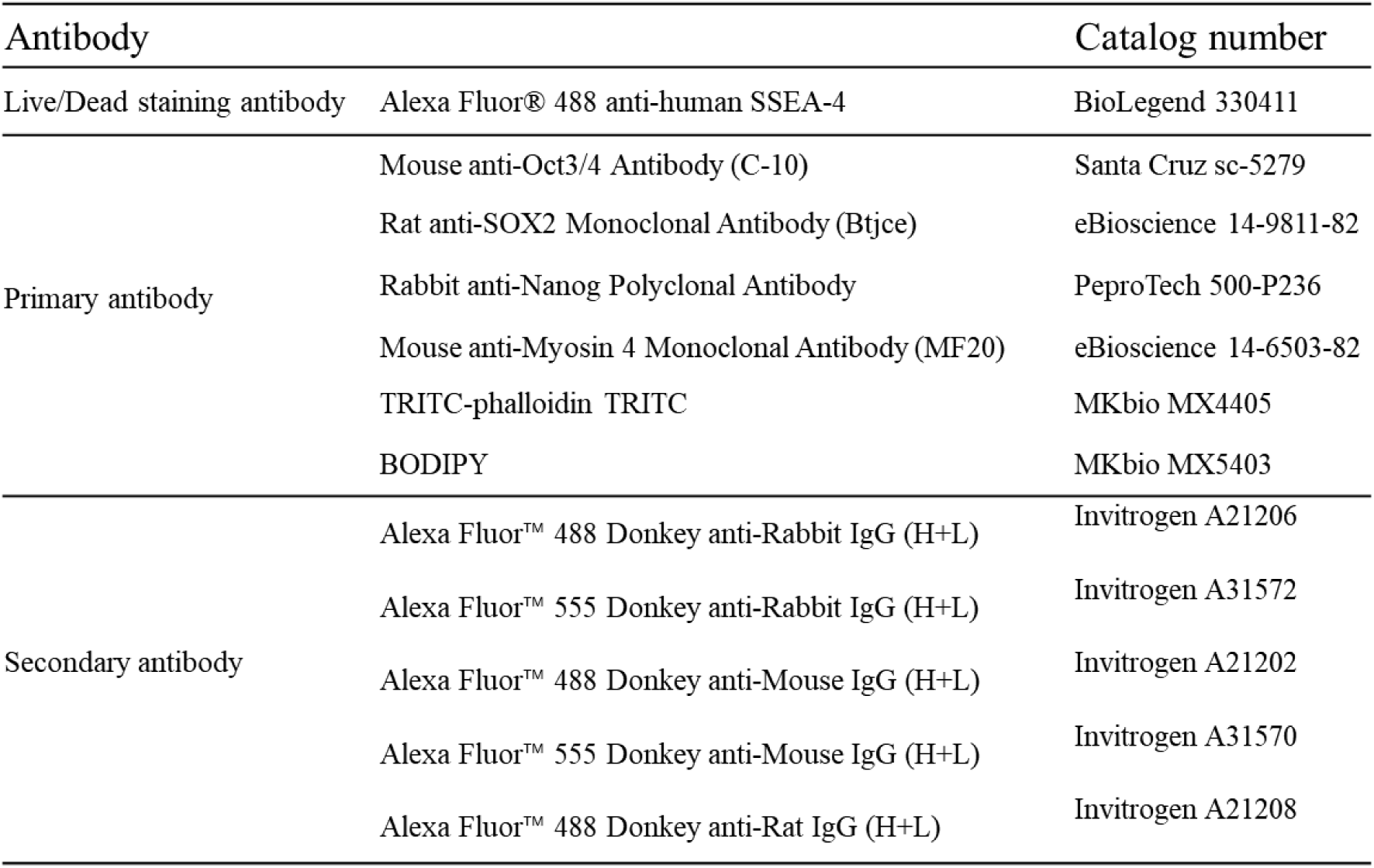
Detailed information of antibodies used for staining.

**Table S2.**
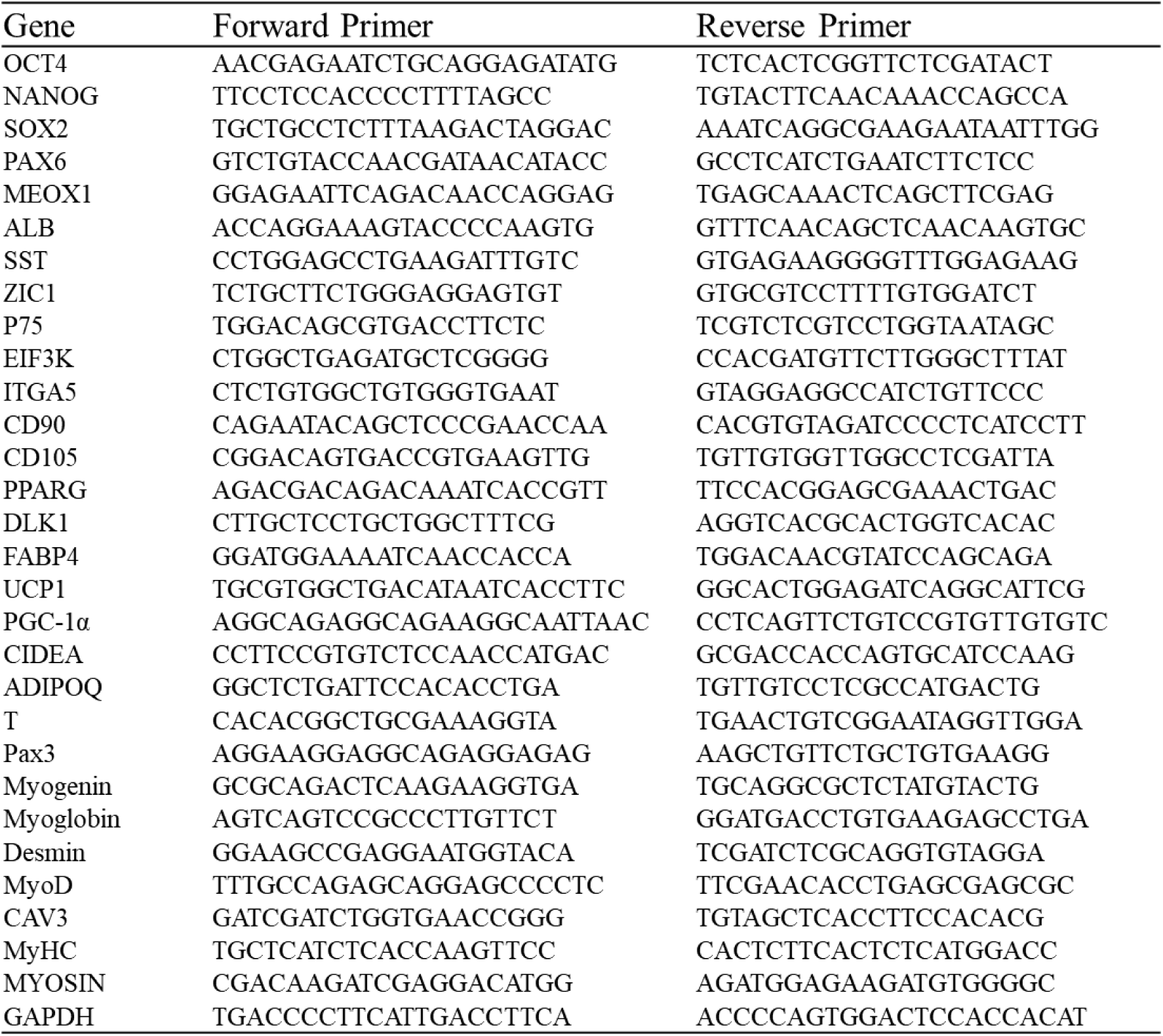
primer sequences used for qRT-PCR.

**Table S3.** The formulation for the production of textured protein scaffolds through the mechanical elongation method.

| Wheat gluten (g) | Soy protein isolate (g) | Water (ml) |
| --- | --- | --- |
| 55 | 32.8 | 8.2 |

**Table S4.** Formulation for scaffold used for cultivated meat.

| Scaffold from table S1 (g) | Wheat gluten (g) | Water (ml) | Corn starch (g) | Beet root (g) |
| --- | --- | --- | --- | --- |
| 13 | 8 | 20 | 6 | 8 |

